# Evolutionary traits of Tick-borne encephalitis virus: Pervasive non-coding RNA structure conservation and molecular epidemiology

**DOI:** 10.1101/2021.12.16.473019

**Authors:** Lena S. Kutschera, Michael T. Wolfinger

**Affiliations:** Department of Theoretical Chemistry, University of Vienna, Währingerstraße 17, 1090 Vienna, Austria; Research Group Bioinformatics and Computational Biology, Faculty of Computer Science, University of Vienna, Währingerstraße 29, 1090 Vienna, Austria

## Abstract

Tick-borne encephalitis virus (TBEV) is the etiological agent of tick-borne encephalitis, an infectious disease of the central nervous system that is often associated with severe sequelae in humans. While TBEV is typically classified into three subtypes, recent evidence suggests a more varied range of TBEV subtypes and lineages that differ substantially in the architecture of their 3’ untranslated region (3’UTR). Building on comparative genomics approaches and thermodynamic modelling, we characterize the TBEV UTR structureome diversity and propose a unified picture of pervasive non-coding RNA (ncRNA) structure conservation. Moreover, we provide an updated phylogeny of TBEV, building on more than 220 publicly available complete genomes, and investigate the molecular epidemiology and phylodynamics with Nextstrain, a web-based visualization framework for real-time pathogen evolution.

## 1 Introduction

Tick-borne encephalitis virus (TBEV) is a zoonotic RNA virus in the genus *Flavivirus,* family *Flaviviridae.* It is the etiological agent of tick-borne encephalitis (TBE), an infection of the central nervous system that is considered the most common tick-transmitted disease in Eurasia [1], where it occurs in risk or endemic areas that are also referred to as foci[2]. TBEV is transmitted between haematophagous ticks as vectors and vertebrate hosts. Typical reservoir hosts include wild-living animals such as small rodent species. Large vertebrate species, including humans and ungulates like goats, cows, sheep, swine, and deer can become infected but appear not to be competent of transmitting the virus back to ticks [3]. While serological evidence suggest that the majority of human infections is either asymptomatic or subclinical, TBEV is a neurotropic virus that can cause a wide range of life-threatening clinical manifestations comprising febrile, meningeal, meningoencephalitic, poliomyelitic, polyradiculoneuritic and chronic forms (reviewed in [4]), as well as hemorrhagic syndrome [5].

### 1.1 Flavivirus genome organization

TBEV belongs to the group of tick-borne flaviviruses (TBFVs), which, together with mosquito-borne flaviviruses (MBFVs) and no-known-vector flaviviruses (NKVs) encompass the vertebrate-infecting flaviviruses. Contrary, insect-specific flaviviruses (ISFVs) only replicate in mosquitoes [6]. Flaviviruses are enveloped, single stranded (+)-sense viruses that contain a non-segmented, 5’-capped, non-polyadenylated RNA of approximately 11kB length. The genomic RNA (gRNA) encodes a single open reading frame (ORF) that is flanked by highly structured untranslated regions (UTRs) of variable length [7, 8]. Both UTRs are crucially involved in regulating processes that control different aspects of the virus life cycle, including virus replication, genome cyclisation and packaging, and immune response [9, 10, 11].

Common architectural traits of flavivirus 3’UTRs comprise autonomous RNA structure formation of distinct domains and the presence of evolutionary conserved RNA elements with specific functional associations. A hallmark of flavivirus biology is their ability to actively dysregulate the host mRNA turnover machinery by stalling endogenous exoribonucleases [12] at structurally well-defined RNAs in the viral 3’UTR [13]. Homologs of these so-called exoribonuclease-resistant RNAs (xrRNAs) are typically found in one or two copies throughout all ecological groups of the genus Flavivirus [14]. Another element that is characteristic of flavivirus 3’UTRs is the long 3’-terminal stem-loop (3’SL) structure, which is is involved in genome cyclisation and panhandle formation during virus replication [15]. As this element is indispensable for the virus life cycle, absence of a 3’SL homolog in sequence data is indicative of incomplete sequencing or truncation. Several other structured RNAs of known and unknown function are found in flavivirus 3’UTRs, including dumbbell (DB) elements and various stem-loop (SL) structures that are either conserved throughout the entire genus or selectively found in particular virus species [16].

### 1.2 TBEV diversity correlates with pathogenicity

TBEV has traditionally been classified into three subtypes, European (TBEV-Eur), Siberian (TBEV-Sib) and Far Eastern (TBEV-FE) [17], based on serology, phylogenetic reasoning and geographic location of early samples. Increased sampling over the previous decades resulted in the availability of more isolates that did not cluster with the established subtypes (Figure 1). As such, additional lineages and provisional subtypes have been reported. These include strains that have been isolated in Eastern Siberia and that have accordingly been designated as “Baikalean” isolates, i.e. genotype 4 (TBEV-Bkl-1), represented by the single strain 178-79 [18], and genotype 5 (TBEV-Bkl-2), encompassing strain 886-84 and related East-Siberian isolates [19, 20, 21]. Moreover, isolates found in wild rodents in the Quinghai-Tibet Plateau, China, constitute a Himalayan TBEV subtype (TBEV-Him) [22]. The Siberian subtype, with prototype strain Aina, is considered the most abundant TBEV subtype [23], and is subdivided into five lineages: Zausaev, Vasilchenko, Baltic, Bosnia, and Obskaya [24, 25, 26, 27], with Obsakya also being referenced as TBEV-Ob subtype in the literature [28]. The Far-Eastern subtype encompasses three clades that have been referred to as Cluster I-III [29]. Each cluster has a prototype strain, i.e. Oshima (Cluster I), Sofijn (Cluster II) and Senzhang (Cluster III). The European subtype comprises lineages containing the Neudoerfl and Hypr stains, as well as a recently described Western European lineage that has been found in the Netherlands [30]. The Austrian strain N5-17, isolated from a chamois, shares ancestral roots with the other European clades. Notably, the European TBEV isolates are phylogenetically closer to Louping ill virus (LIV), which is present in the British Isles [31], than to the Eastern TBEV subtypes.

**Figure 1:**
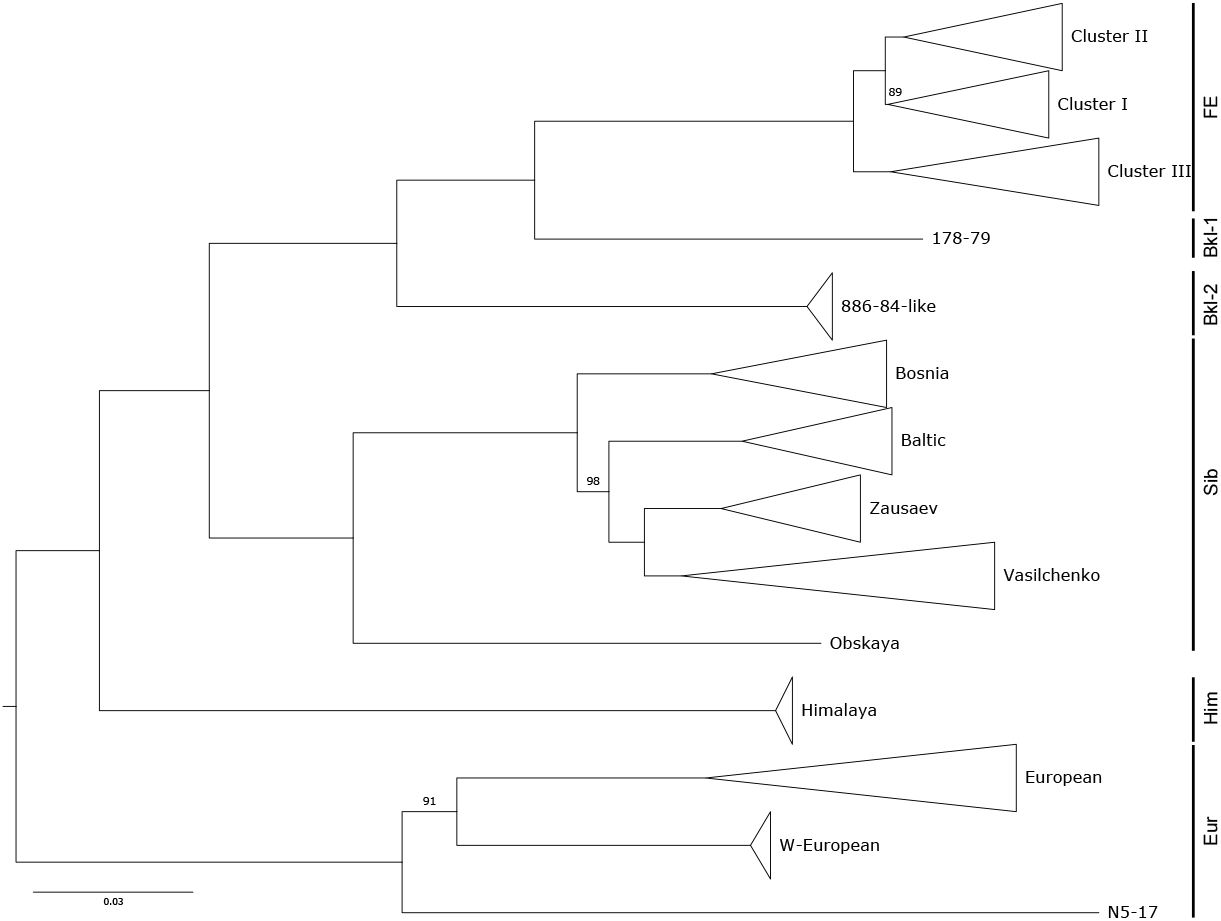
Overview phylogeny of TBEV, depicting the topological arrangement of subtypes TBEV-FE, TBEV-Sib, and TBEV-Eur (all highlighted *sensu strictu* here), as well as novel lineages that do not cluster with these subtypes. Strain 178-79 and the Baikalean lineage share ancestral roots with TBEV-FE. The Himalayan lineage is located ancestral to the clades encompassing TBEV-FE and TBEV-Sib subtypes. The European TBEV strains form a separate clade, with the recently detected Western European and N5-17 lineages being clearly separated from the established TBEV-Eur subtype. The midpoint-rooted maximum likelihood tree is based on 256 complete TBEV genomes. Bootstrap values are shown for branches with a support lower than 100.

In a recent study, the genomic diversity of established an provisional TBEV subtypes has been assessed and classification of TBEV strains into seven subtypes has been proposed based on a 10% cutoff at nucleotide-level diversity [28]. Despite their antigenic similarity, these viruses differ not only in their phylogeography, but also in virulence and pathogenicity. The Far Eastern subtype is commonly considered the most pathogenic TBEV variant with the highest number of reported cases of severe central nervous system (CNS) disease. Case fatality rates of TBEV-FE are typically in the range of 5%-35% [32], with extreme numbers reaching up to 60% [4]. Contrary, the Siberian subtype typically results in less severe disease with a fatality rate between 1% and 3% [4], but is more often associated with chronic TBE [2]. Albeit case fatality at <2% is least for the European subtype [33], disease induced by TBEV-Eur infection is typically biphasic with a viraemic phase associated with fever and myalgia, followed by neurological manifestations of different severity during the second phase, which occurs in 20%-30% of all patients [4].

### 1.3 Functional RNAs in TBEV non-coding regions

The fundamental biological and biochemical traits that govern TBEV pathogenicity are only beginning to be understood. Besides several subtype-specific differences at the amino acid level [29, 34], the non-coding portions of the TBEV genome are critical determinants of virulence. The 5’UTR is approximately 130nt long and contains evolutionarily conserved RNA struc-tures that are required for replication through cyclisation and panhandle formation. Recently, a cis-acting RNA element in the 5’UTR of TBEV has been identified that mediates neurovirulence by hijacking the host mRNA transport system, thus allowing TBEV gRNA to be transported from the cell body to dendrites of neurons, where it replicates locally [35, 36].

The TBEV 3’UTR is longer than the 5’UTR and contains RNA elements that are involved in mediating immune escape and pathogenicity. It is characterized by a varied architecture of conserved RNA elements that has been associated with differential pathogenicity of particular subtypes [29, 37]. Although a few TBEV-FE isolates exist in public databases that appear to have extremely shortened 3’UTRs of only approximately 50 nt length, typical 3’UTRs range from 350 nt to 760 nt and comprise a 5’-terminal variable region, located immediately downstream of the stop codon, and a 3’-terminal core domain. Earlier comparative studies that have assessed the heterogeneity of these genomic regions at the sequence level have proposed various directed repeat sequences [38, 39]. Considering RNA structure formation [7] allowed for a more fine-grained description and detailed comparative analysis of evolutionary conserved RNA elements with other tick-borne flaviviruses [14].

An interesting aspect relates to the observation that strains of a particular subtype that have been isolated from ticks or rodents, respectively, are often markedly different from human isolates, particularly in their 3’UTRs. This might be the result of selective pressure on the TBEV 3’UTR during adaptation from tick vector to mammalian host [40]. Likewise, spontaneous deletions of parts of the variable region have been observed after repeated passage [41, 37]. This is complicated by the claim that loss of 3’UTR elements in infected mammalian cells is not quantitative, leading to a scenario that different portions of the viral population have different 3’UTR architectures, each having different replicative fitness when transmitted to ticks [29]. Given that strains resulting in different disease manifestations, ranging from subclinical to encephalitic forms, cluster within the same clades or subclades of the TBEV phylogeny [29], the question of functional association and impact on clinical phenotype of different 3’UTR elements is emerging. There are contradictory reports in the literature regarding the influence of deletions in the 3’UTR variable region on the TBEV phenotype. While earlier works proposed no role of the variable region in virus replication and virulence [41, 42], recent studies suggest increased virulence induced by partial deletions and polyA insertions in mouse models [43, 44].

In this study, we investigate the genetic diversity and phylogeographic spread of TBEV, based on a comprehensive set of more than 220 genomes that encompasses all known and provisional subtypes. By considering also strains that do not cluster with the established subtypes, we provide a unified picture of functional RNA conservation in TBEV 3’UTRs. Moreover, we contextualize genomic and epidemiological data through Nextstrain, providing an updated online resource to study the molecular evolution of TBEV.

## 2 Methods

### 2.1 Taxon and metadata collection

Viral genome and annotation data were downloaded from the public National Center for Biotechnology Information (NCBI) Genbank database [45] on 14 January 2021. We obtained 256 TBEV genomes, containing at least full length coding sequences. 221 sequences contained partial or full length 3’UTRs. We extracted metadata for these strains, including location and date of collection, subtype and host species from the Genbank record or from the literature. More than two dozen sequences that have been identified as vaccine strains, highly cell-passaged, or that were the result of cloning, mutants, or duplicates, were filtered out and excluded from downstream analyses. Strain identifiers which were not available through online NCBI resources, but obtained by literature research, were added manually.

Countries were assigned to regions according to the M49 Standard of the United Nations Statistic Division (https://unstats.un.org/unsd/methodology/m49/). A categorization of host species was included, which assigns the species of which a particular strain was isolated from to a category according to their role in viral transmission. These categories were vector (mites, ticks), natural host (rodentia, insectivora, aves), non-human dead-end host (large mammalia) or human.

### 2.2 TBEV subtype and lineage classification

To assign each strain in our dataset to a particular TBEV subtype or lineage, we aligned 221 (near) full-length TBEV genomes with `MAFFT` v7.453 [46] and inferred a maximum likelihood (ML) phylogeny with `iq-tree` v1.6.12 [47] (Figure 1), employing 1000 ultrafast bootstrap replicates [48]. Strains with initially unknown subtype association were assigned to subtypes via distinguishing subtype-or lineage-specific monophyletic clades in the ML tree. Upon classification of strains, we used the multiple nucleotide sequence alignment (MSA) to identify all unique 3’UTR variants within each subtype and lineage, and selected a representative strain for each instance (28 strains, listed in Table S1).

### 2.3 Characterization of RNA consensus secondary structures

Detection of evolutionary RNA structure conservation builds on the idea of finding homologous RNAs in phylogenetically narrow taxa via RNA family models [49]. Here, we used `infernal` [50] covariance models (CMs) and employed an iterative workflow that has been used recently to characterize evolutionary conserved RNAs in MBFVs [16]. Starting from a set of TBFV CMs described in [14], we performed an initial screen of TBEV 3’UTRs. Genomic regions that were not covered by the initial CMs were then realigned with `locARNA` v2.0.0RC8 [51] and RNA consensus structures were predicted with `RNAalifold` and `RNALalifold` from the `ViennaRNA` Package v2.4.18 [52].

### 2.4 Nextstrain Phylogeography

The workflow management system `Snakemake` v5.32.1 [53] was used to build a pipeline for rapid and reproducible deployment of Nextstrain [54] datasets. `augur` v11.0.0 [55] was used for tracking evolution from nucleotide sequence data, incorporating `MAFFT` v7.453 [46], `iq-tree` v1.6.12 [47], and `TreeTime` v0.8.4 [56], for MSA, ML phylogeny, and timed tree inference, respectively, resulting in a JSON file for Nextstrain visualization. Values for the rate of evolution were chosen according to the molecular clock rate that has been previously described as 5.96e-5 with standard deviation of 6.6e-6 substitutions per site and year [57].

### 2.5 Data availability

Stockholm MSA files of the conserved RNA structures are available at the viRNA GitHub repository (https://github.com/mtw/viRNA). The TBEV Nextstrain build is available at https://nextstrain.org/groups/ViennaRNA/TBEVnext/. Upon display of the TBEV build in a web browser, all data can be downloaded by scrolling down to the bottom of the page and clicking on the “Download Data” link.

## 3 Results

To obtain an updated view of TBEV subtype diversity, we compared publicly available genome data from a structural and molecular epidemiological perspective. For the structural analysis, we assessed the architectural organization of TBEV 5’UTRs and 3’UTRs, the latter constituting the most divergent part of the genome. We performed a comparative genomics screen that builds on the concept of predicting consensus structures, i.e. RNA secondary structure that can be formed by all sequences under consideration. We propose a conserved 5’UTR organization, as well as the existence of eight distinct, evolutionarily conserved RNA elements in the 3’UTR of different TBEV subtypes. The volatile arrangement of these non-coding RNAs (ncRNAs) at inter- and intra-subtype levels accounts for the diverse TBEV 3’UTR architectures observed in nature.

### 3.1 TBEV 5’UTRs are structurally conserved

The TBEV 5’UTR has a canonical length of 132 nt throughout all subtypes. Despite conservation of 5’UTR length, there is considerable sequence variability that accounts for average sequence identity levels between 89% and 95% for TBEV-Sib and TBEV-FE, respectively [58]. This heterogeneity at the primary sequence level, however, does not translate into structural diversity. Consensus structure prediction yielded a clear picture of uniform 5’UTR organization throughout all TBEV subtypes and lineages: The first element of the TBEV genome is a variant of the Y-shaped stem-loop A (SLA) structure, which has been associated with panhandle formation and recruitment of the viral RNA-dependent RNA polymerase during virus replication [59]. Downstream of the SLA element, a short hairpin (CSA) is located, which is followed by another hairpin (CSB) that overlaps the AUG start codon of the capsid protein coding region. Figure 2 shows the annotated 5’UTR of the TBEV reference strain Neudoerfl. Among the three 5’UTR-associated RNA structural elements, SLA and CSB exhibit covariation patterns, while CSA is highly sequence-conserved (Figure S1).

**Figure 2:**
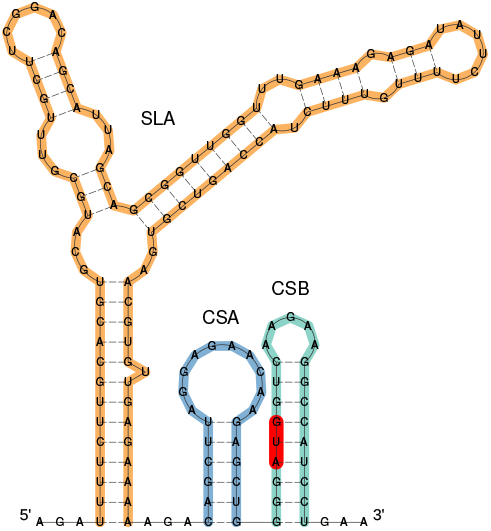
Secondary structure prediction of the 5’-terminal 159 nt of TBEV strain Neudoerfl (NC_001672.1), comprising the 5’UTR and the distal portion of the capsid protein coding region. The canonical 5’UTR organization of TBEV encompasses a Y-shaped SLA element (orange), as well as the hairpin loops CSA (blue), and CSB (green). The start codon is highlighted in red.

### 3.2 TBEV 3’UTRs comprise up to eight distinct RNA elements

Starting from nucleotide multiple sequence alignments of 221 TBEV genomes that covered all known subtypes and lineages, we set out to construct consensus RNA structures of conserved elements. Following up and building on previous work [14], we identified seven distinguishable families of conserved RNA structures that are found as single, double, or triple copies in a variable arrangement in TBEV strains. Together with the poly-A tract that is found in some TBEV-Eur strains, we characterized eight unique ncRNA elements in TBEV 3’UTRs. Inconsistencies with the naming schema of structured TBEV 3’UTR elements in the literature have lead us to introduce here a pragmatic naming system for conserved ncRNA elements.

The structured elements encompass four distinct conserved stem-loop (CSL) elements of unknown function, labeled CSL1 through CSL4, starting from the most 3’-terminal elements. Predicted consensus secondary structures of these elements are shown in Figs. S2 and S3. CSL1, CSL2, and CSL4 are short stem-loop elements that are found in variable copy numbers in different TBEV strains. We have previously shown evolutionary conservation of CSL2 in other tick-borne flaviviruses, where this element was referred to T.SL6 [14]. CSL4, which has a canonical length of approximately 30 nt, has a consensus structure that contains multiple gap characters in the hairpin loop. This is due to the insertion of a poly-A region of variable length in the apical loop of one CSL4 copy in some TBEV-Eur strains. Contrary, CSL3 is markedly longer than the other CSL elements and is interspersed with several interior loops. Comparison of covariance models with CMCompare revealed that CSL3 is not structurally homologous to SL-III elements that were found in various mosquito-borne flaviviruses [16].

Besides the CSL elements, TBEV 3’UTRs contain previously described ncRNAs with known functional associations, such as three-way junction elements that form exoribonuclease-resistant RNAs (xrRNAs). These elements, which are ubiquitous in the viral world [60, 61], provide quantitative protection of downstream nucleotides against degradation by endogenous exoribonucleases and have been structurally and mechanistically characterized in different tick-borne flaviviruses [13, 62]. A Y-shaped structure (Y1), together with the the 3’-terminal part of TBEV 3’UTRs, encompassing a small hairpin loop and a long stem-loop element, labeled 3’SL here, comprise the promoter element, which is essential for virus replication. Screening for structural homology of the above elements in other, i.e. non tick-borne flaviviruses corroborated earlier knowledge that xrRNA and 3’SL elements are ubiquitously conserved throughout the genus *Flavivirus.* While the CSL elements do not appear to be universally conserved outside tick-borne flaviviruses, a structurally homologous Y1 element was found in the 3’UTR of Modoc virus (MODV), a tick-borne related no-known vector flavivirus.

A structural annotation of TBEV 3’UTRs is shown in Fig. 3, which highlights the differences in architectural organization among and within TBEV subtypes. Selection of strains featured in Fig. 3 was based on subtype diversity, i.e. coverage of all established and prospective subtypes, as well as within-subtype variability of 3’UTR organization. We included at least one representative isolate of each 3’UTR variant that has been present in our data set. Full-length 3’UTRs (where available) are plotted to scale in Figure 3, with colored boxes representing evolutionary conserved RNA elements. Homologous elements are shown in the same color for each strain.

**Figure 3:**
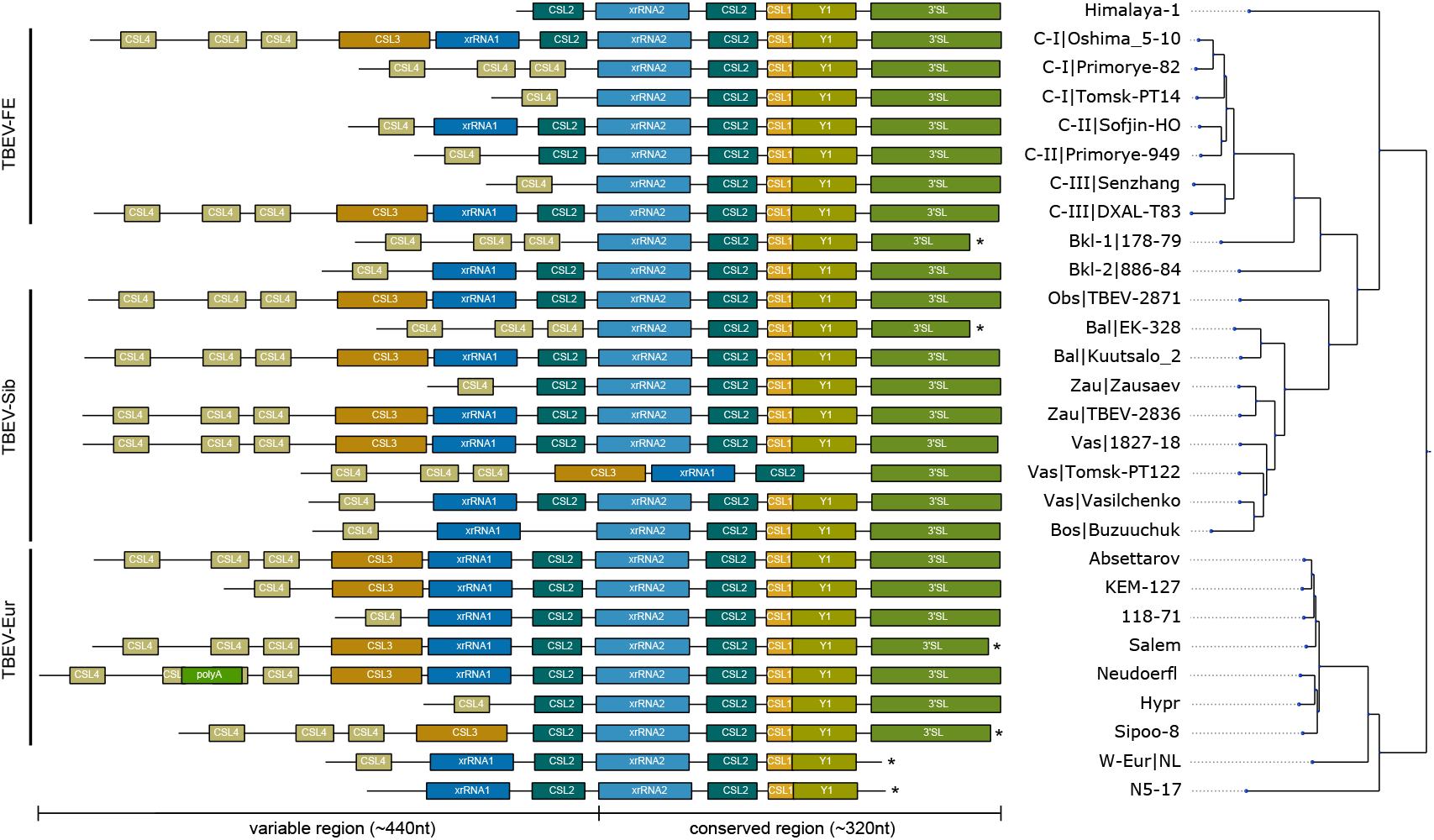
Annotated 3’UTRs of representative TBEV strains. Strains were selected to cover the complete 3’UTR diversity found in publicly available genome data. 3’UTR schemes are plotted to scale, highlighting the variable length and architectural organization of individual strains. Colored boxes represent evolutionary conserved, structured RNAs. Truncated sequences from incomplete sequencing are marked by an asterisk. The phylogenetic tree on the right has been computed from full genome nucleotide multiple sequence alignments of 28 TBEV strains. Identifiers show strain name, and lineage association where available, i.e. TBEV-FE Clusters I/II/III (C-I/C-II/C-III), the two Baikalean subtypes TBEV-Bkl-1 (Bkl-1), and TBEV-Bkl-2 (Bkl-2), and TBEV-Sib lineages Baltic (Bal), Zausaev (Zau), Vasilchenko (Vas), Bosnia (Bos), Obskaya (Obs). Accession numbers and 3’UTR lengths are listed in Table S1.

Our comparative analysis of TBEV 3’UTRs suggests a common pattern of structural conservation, revealing that both core and variable regions contain specific RNA elements that are characteristic of their location within the 3’UTR. This is particularly marked in the core region, which contains five highly conserved structured RNA elements within approximately 320 nt. These include an exoribonuclease-resistant RNA (xrRNA2), CSL2, CSL1, Y1, and the 3’-terminal long stem-loop structure (3’SL). Consensus secondary structures of core region elements are shown in Fig. S2. The well-conserved architecture of potentially functional RNAs in the core region is observed in all strains considered here, except Tomsk-PT122 (TBEV-Sib).

The variable region ranges between approximately 80 nt (Tomsk-PT-14) and 440 nt (Neudoerf I), and contains up to five different types of RNA elements, i.e. CSL4, poly-A, CSL3, xrRNA1, and CSL2. Consensus secondary structures are shown in Fig. S3. Some of these RNAs occur in multiple copies at particular loci. Most strains contain between one and three copies of CSL4 at the 5’-terminal portion of their 3’UTRs. There are only two exceptions to this observation within our set of strains, i.e. Himalaya-1 and N5-17. The internal poly-A region observed in some European strains is inserted into the hairpin loop of the second CSL4 copy. While a variable number of CSL4 homologs is found in almost all strains, a long variant of the remaining portion of the variable region, containing CSL3, an exoribonuclease-resistant RNA (xrRNA1), and CSL2, is only conserved in some strains. These long 3’UTR variants are found in all subtypes, e.g. Oshima5-10 (TBEV-FE), 1827-18 (TBEV-Sib), and Absettarov (TBEV-Eur). The structured elements CSL3, xrRNA1 and CSL2 in the variable region appear to be dispensable for virus replication, as other strains either lack a CSL3 element (e.g. Sofjin-HO or Vasilchenko), CSL3 and xrRNA1 (e.g. Hypr or Zausaev), or CSL3, xrRNA1 and CSL2 (e.g. Senzhang or Tomsk PT14).

To illustrate the differences in 3’UTR architectures due to alternative layout of the variable region, we show secondary structure plots of four complete 3’UTRs in Fig. 4. With all strains exhibiting a common core region architecture, we depict representative strains of the European subtype (Neudoerfl, panel A) and the Obskaya lineage of the Siberian subtype (TBEV-2871, panel B), that both exhibit long variants of the variable region. These comprise three CSL4 copies, CSL3, xrRNA1 and one CSL2 element. Insertion of a poly-A stretch results in the formation of a large hairpin loop in the second CSL4 copy of the Neudoerfl strain. The 886-84 strain, representing the Baikalean subtypes, (panel C) has a considerably shorter variable region that contains a single CSL4 copy, xrRNA1, and CSL2, with additional CSL4 structures and CSL3 missing. The Shenzhang strain of the Far Eastern subtype (depicted in panel D) has the shortest 3’UTR in our data set, and contains just one CSL4 element in the variable region.

**Figure 4:**
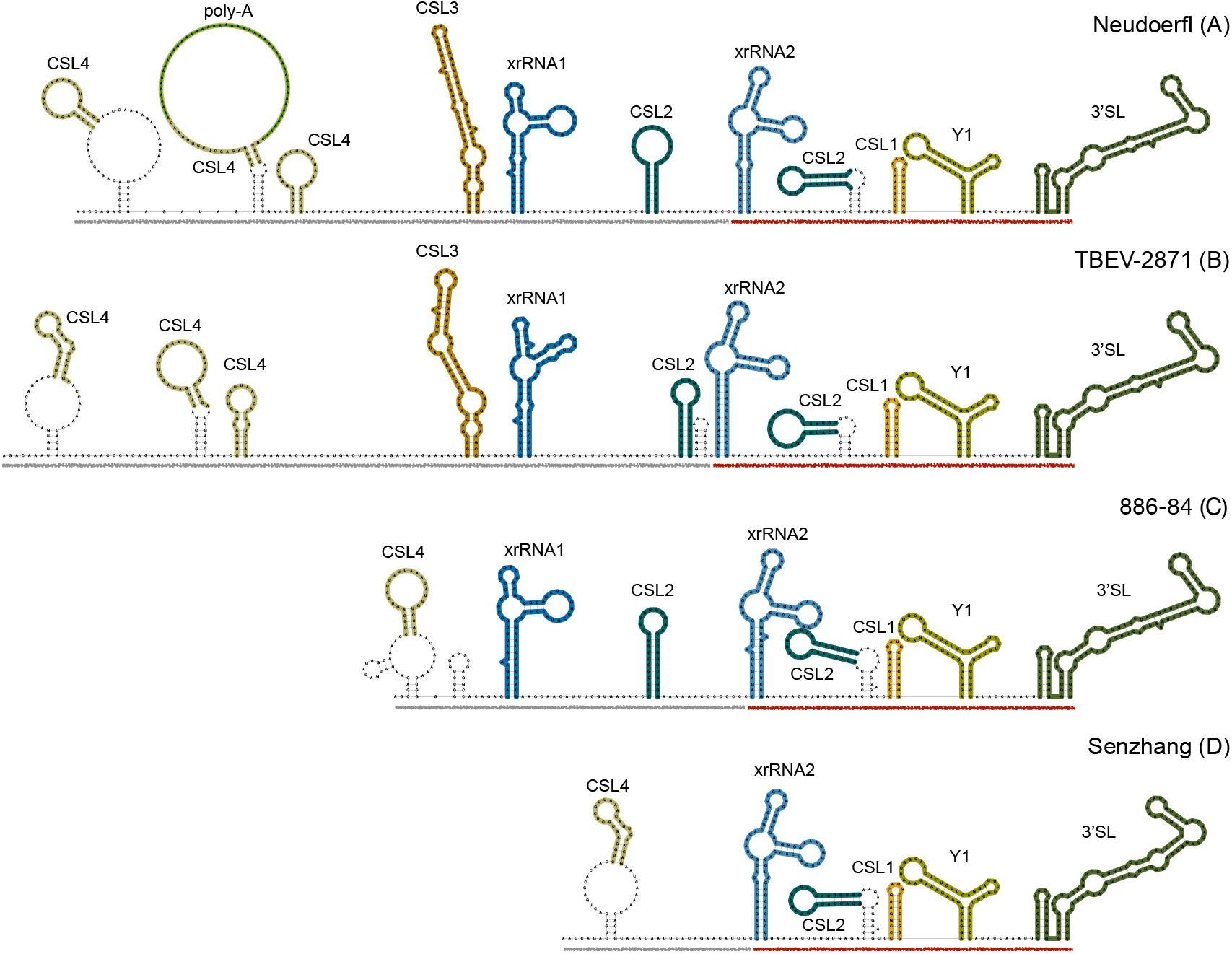
Diverging 3’UTR architecture in four representative TBEV strains, Neudoerfl (A), TBEV-2871 (B), 886-84 (C), and Senzhang (D). Core and variable regions are underlined in red and grey, respectively. Structural homologs of evolutionarily conserved RNA elements are depicted in the same color for all strains (matching the colors used in Fig. 3). Structural components that are not found in all homologs, such as closing stems of some CSL4 and CSL2 elements, or additional short stem loops that are only predicted in particular strains, are not considered conserved.

Some of the recently described strains have a 3’UTR organization that is also found in other lineages. The Western European strain NL and N5-17, while both being truncated at the 3’ end, show similar patterns e.g. to strain 886-84. Contrary, other strains exhibit alternative 3’UTR architectures that are manifested by unique organizational patterns in the variable region that are not observed elsewhere. For example, the 3’UTR of Himalaya-1 does not have a CSL4 element at the 5’-end, but begins with a CSL2 structure. The Buzuuchuk strain, representing the Bosnian lineage of TBEV-Sib, does not have a CSL2 element in the variable region. Sipoo-8, collected in Finland, lacks the exoribonuclease-resistant RNA element in the variable region, although it has three CSL4 elements and CSL3. The most unconventional 3’UTR organization is found in Tomsk-PT122 (Vasilchenko lineage, TBEV-Sib), which is the only example of a strain that exhibits deletions the core region. Tomsk-PT122 is predicted to contain the long version of the variable region described above, while xrRNA2, CSL2, CSL1 and Y elements are are not found in the core region.

### 3.3 TBEVnext: TBEV molecular epidemiology

To get a comprehensive picture of TBEV phylogeography, considering all publicly available isolates, we sought to to investigate the molecular epidemiology and phylodynamics of TBEV. To this end, we compiled an interactive visualization of the global TBEV spread in Nextstrain, termed TBEVnext, which is publicly available at https://nextstrain.org/groups/ViennaRNA/TBEVnext. At the time of writing, TBEvnext v1.0 has been made available with 225 TBEV strains, encompassing all subtypes and lineages.

The overall topology of our TBEV phylogeny (Figure 5a) is in agreement with the literature, featuring Eastern (TBEV-FE, TBEV-Sib), and Western (TBEV-Eur) types as major clades [63]. Strains that are not assigned to one of the larger clades are 178-79 (TBEV-Bkl-1), which is located ancestral to the TBEV-FE clade, and N5-17, which appears ancestral to all other European clades. The East-Siberian/Baikalean/886-84-like (TBEV-Bkl-2) lineage, as well as the Western-European and Himalayan lineages are smaller monophyletic groups that comprise between two and eight isolates. TBEV-Bkl-2 clusters with the Far-Eastern clade, Western European isolates cluster with TBEV-EUR, and the Himalayan isolates share ancestral roots with the clade that comprises all Eastern TBEV isolates. Divergence of the different TBEV subtypes and lineages is exposed in the TBEVnext dataset through an unrooted phylogenetic tree (Figure 5b).

**Figure 5:**
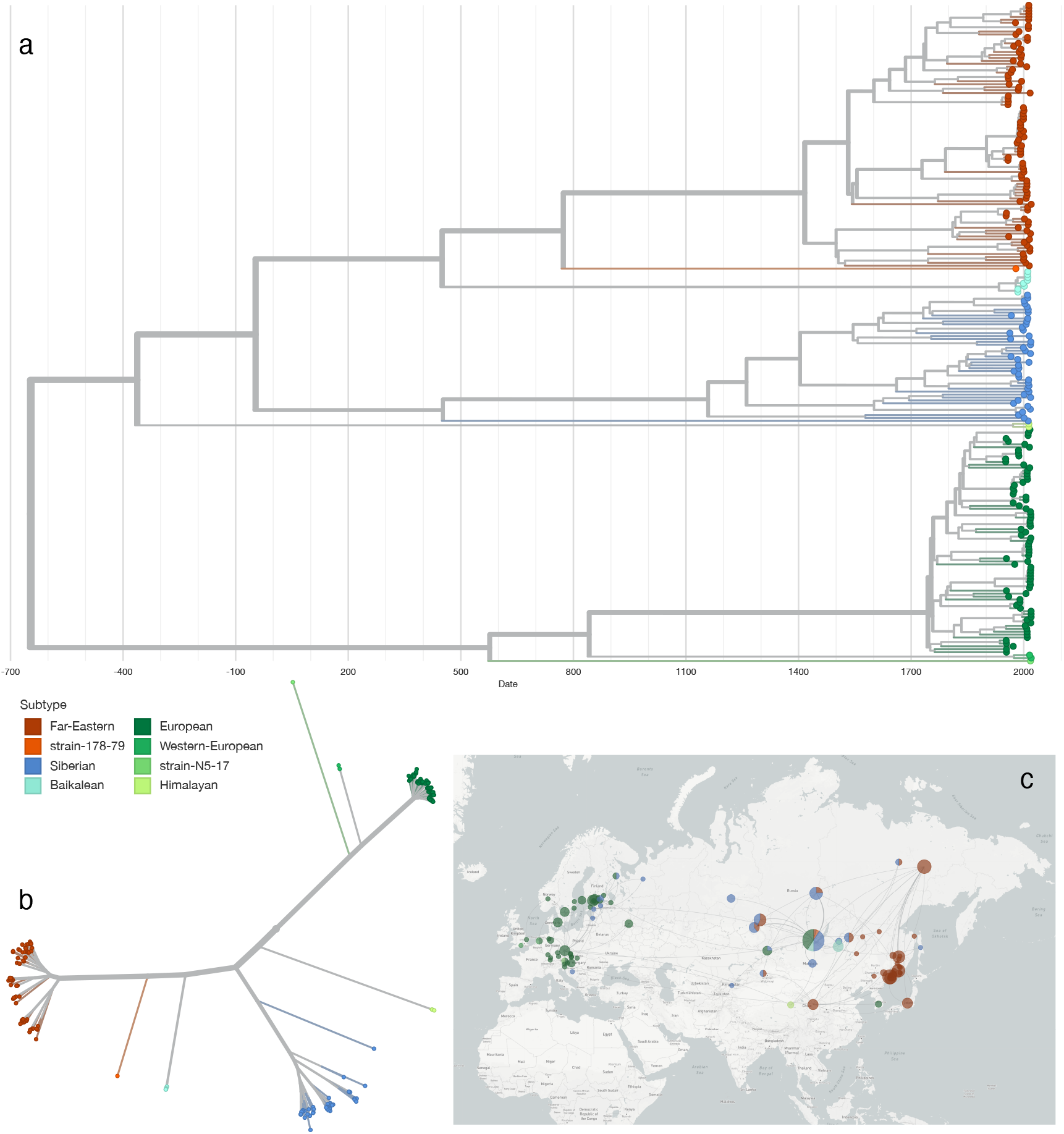
**a** Visualization of the TBEV phylodynamics in Nextstrain. **b** Unrooted phylogenetic tree, exposing the divergence of TBEV subtypes and novel lineages. **c** Geographic spread of TBEV across Eurasia.

The Nextstrain framework is particularly useful for studying phylogeographic traits. As such, we established a fine-grained geographic location labeling of strains that had relevant location information (i.e. place of isolation) in their metadata. While an exact place of isolation could not be inferred for all strains, sub-national geo-locations are available for many isolates in TBEVnext. This is particularly relevant for studying the spread of different subtypes across geographically extended regions, such as presence of various subtypes in the Russian Federation. An example is Irkutsk Oblast, which shows an accumulation of different subtypes: Western, East-Siberian, Siberian, Far-Eastern and strain 178-79 were collected in this region.

The TBEV Nextstrain dataset features a timed tree, which provides estimates for the time of the most recent common ancestor (TMRCA) associated with each clade (Table 1). The timed tree has been inferred from a ML tree with treetime, assuming a constant evolutionary rate throughout the entire tree. Diverging nucleotide substitution rates have been proposed for different TBEV subtypes [57, 64, 65], hence our numbers should be seen as a rough estimate.

**Table 1:**
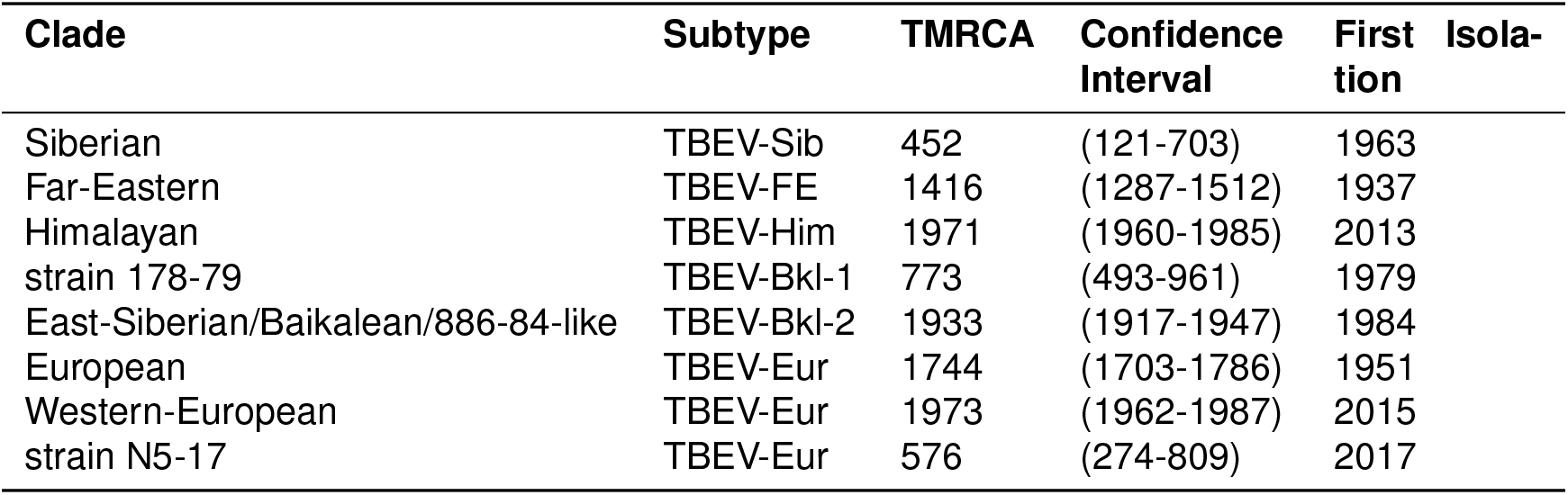
TMRCA estimates of distinct clades, and associated subtypes of the TBEV phylogeny, extracted from the TBEV Nextstrain dataset. All dates are given as year CE.

Animation of historic dispersal events across Eurasia within Nextstrain suggests that TBEV originated in Central Russia almost 2700 years ago, and subsequently moved both eastwards, forming the TBEV-Sib and TBEV-FE lineages, and westwards into Europe. While this observation is in agreement with earlier reports [63], the deep splits of the Western types with inferred TMRCAs approximately 1000-1500 years ago raises the question whether TBEV has arrived in Central Europe earlier than previously reported.

One of Nextstrain’s strengths is the possibility to augment the dataset with custom traits, in addition to default attributes like date and place of collection. Adopting this feature, we maintain in TBEVnext a mapping of species from which a particular strain has been isolated. It has been previously described that different TBEV subtypes are specialized to particular vector species of Ixodidae. The Eastern subtypes (Far-Eastern, Siberian) are typically transmitted via *I. persulcatus,* whereas the Western subtypes (European) are usually transmitted by *I. ricinus* ticks. While this is broadly supported by our data, metadata analysis of 225 full genome isolates also discloses counter-examples: Nine isolates collected from *I. persulcatus* ticks between 1971 and 2009 in Irkutsk Oblast and the Altai region in Russia carried the European subtype, while two *I. ricinus* ticks carried the Siberian subtype. (Data shown in the interactive visualization of TBEVnext.)

## 4 Discussion

Increased sample collection and availability of large amounts of genome data in public databases has stipulated research on flaviviruses in recent years. TBEV, being endemic in large parts of Europe and Asia, has been intensively studied, resulting in numerous publications focusing on the functional associations of the variable 3’UTR region in particular strains and viral (neuro)tropism. Some of these studies reported a variety of sequence repeats in the 3’UTR of tick-borne flaviviruses, based on qualitative comparisons at the nucleotide level. Other studies performed computational modelling of RNA secondary structure at the level of individual sequences, yielding a varied picture of stem-loop structures that makes it difficult to compare structured entities. A unified, reproducible picture of ncRNA structure conservation in different TBEV subtypes has not been available. Here, we set out to obtain an updated view of RNA structure conservation in TBEV 3’UTRs, aiming at capturing a unified picture of the architectural diversity found in different subtypes. To this end, we performed a comparative genomics screen in TBEV 3’UTRs that cover all known subtypes and lineages. The computational approach employed here is based on a thermodynamic model of RNA structure formation, as implemented in the ViennaRNA package.

TBEV 3’UTRs comprise two domains, a core domain at the 3’end of the viral genome, and a variable region upstream of the core domain. The 3’-terminal core domain constitutes the most conserved portion of TBEV 3’UTRs, and harbors five different structured ncRNA elements. Two of these, an exoribonuclease-resistant RNA (xrRNA2) and the terminal 3’ stem-loop element are also known in other ecological groups of flaviviruses and have been associated with mediating virulence and virus replication [16]. The variable region, on the other hand, contains between two and four distinct structural elements. Observed differences in the length of TBEV genomes that belong to different subtypes is primarily due to heterogeneity of the variable region in TBEV 3’UTRs. Here, another xrRNA copy (xrRNA1) constitutes the only element with known functional association, while simple and extended stem-loop structures of unknown function appear in variable copy numbers. Moreover, an internal poly-A tract is found in several European isolates [66].

TBEV fits well into the picture of selective conservation of RNA structure in the viral 3’UTR, with core and variable regions containing functional RNA elements, i.e. xrRNAs, that are also found in other arboviruses. Unlike other viruses that exhibit strictly lineage-specific patterns of 3’UTR architecture, such as Chikungunya virus [67, 68], TBEV shows a large degree of 3’UTR variability within particular subtypes and lineages. Our data highlight that there is no common 3’UTR architecture associated with certain TBEV subtypes besides the core region. On the other side, we show that the TBEV 3’ architectural diversity is realized by combination of a restricted pool of eight structural and one non-structural elements, suggesting that the observed 3’UTR variability is driven by structural, rather than sequence conservation.

Beyond studying evolutionary traits of TBEV at the level of RNA structure conservation, we address the issue also from the perspective of molecular epidemiology. To this end, we provide TBEVnext, a Nextstrain dataset that reveals spatiotemporal and phylodynamic characteristics of the global TBEV spread. Our dataset comprises 225 complete genomes from Eurasia that have been sampled between 1951 and 2019, thereby representing the most comprehensive view of TBEV phylogeographics so far.

In addition to the three major TBEV subtypes TBEV-Eur, TBEV-Sib, and TBEV-Eur, TBEVnext encompasses provisional subtypes like TBEV-Bkl1, TBEV-Bkl-2, and TBEV-Him. The Obskaya lineage of TBEV-Sib, deemed an independent subtype in the recent literature, is considered part of the Siberian clade here, while strain N5-17 is considered part of TBEV-Eur. Analysis of the geographic dispersion of these clades in TBEVnext immediately reveals that they are not specific for particular regions of the world. Lack of an unique association between TBEV subtypes (clades) and geographic occurrence is particularly evident in Irkutsk Oblast, where different strains of Siberian, Far Eastern, Western, and Baikalean subtypes have been isolated. Likewise, presence of Western strains in the Republic of Korea strains has been reported [69].

TBEVnext features a timed phylogenetic tree that has been inferred from a maximum-likelihood phylogeny under the assumption of a constant evolutionary rate, effectively neglecting the possibility of divergent rates in different parts of the tree. Therefore, our timed tree should be seen as a proxy that is in agreement with proposed TBEV tree topologies and divergence patterns. Importantly, the order of TMRCAs of different clades is in overall agreement with published data that were based on computationally more demanding Bayesian approaches [21].

Availability of global vector/host associations of many strains in TBEVnext yields a comprehensive view on ecological aspects of the global virus spread, and makes unique properties of strains accessible.

In summary, we provide here an updated picture of different evolutionary aspects of tick-borne encephalitis virus. Our data expose patterns of pervasive conservation of individual RNA elements in TBEV 3’UTRs and provide an interactive view of TBEV molecular epidemiology that yields novel insight into the global virus spread.

## 5 Acknowledgements

We thank Gabor Erdo Mate for fruifutful discussions during the preparation of the manuscript.

