## Supplementary data for "Evolutionary traits of Tick-borne encephalitis virus: Pervasive non-coding RNA structure conservation and molecular epidemiology"

**Table S1:** Representative strains used in this study.

| Accession | Subtype | Strain | Note | 3'UTR length |
|---|---|---|---|---|
| MG599476.1 | TBEV-Him | Himalaya-1 | - | 384 |
| AB062063.2 | TBEV-FE | Oshima 5-10 | Cluster I | 724 |
| JQ825148.1 | TBEV-FE | Primorye-82 | Cluster I | 509 |
| KJ914682.1 | TBEV-FE | Tomsk-PT14 | Cluster I | 404 |
| AB062064.1 | TBEV-FE | Sofjin-HO | Cluster II | 518 |
| KU761570.1 | TBEV-FE | Primorye-949 | Cluster II | 466 |
| JQ650523.1 | TBEV-FE | Senzhang | Cluster III | 408 |
| MN615728.1 | TBEV-FE | DXAL-T83 | Cluster III | 719 |
| EF469661.1 | TBEV-Bkl-1 | 178-79 | - | 488 |
| KJ633033.1 | TBEV-Bkl-2 | 886-84 | - | 539 |
| MF774565.1 | TBEV-Sib | TBEV-2871 | Obskaya | 725 |
| DQ486861.1 | TBEV-Sib | EK-328 | Baltic | 471 |
| MG589939.1 | TBEV-Sib | Kuutsalo_2 | Baltic | 727 |
| AF527415.1 | TBEV-Sib | Zausaev | Zausaev | 455 |
| MH645618.1 | TBEV-Sib | TBEV-2836 | Zausaev | 730 |
| MN114637.1 | TBEV-Sib | 1827-18 | Vasilchenko | 727 |
| KM019545.1 | TBEV-Sib | Tomsk-PT122 | Vasilchenko | 556 |
| AF069066.1 | TBEV-Sib | Vasilchenko | Vasilchenko | 550 |
| KJ626343.1 | TBEV-Sib | Buzuuchuk | Bosnia | 546 |
| KJ000002.1 | TBEV-Eur | Absettarov | - | 720 |
| MG210947.1 | TBEV-Eur | KEM-127 | - | 617 |
| KY069120.1 | TBEV-Eur | 118-71 | - | 528 |
| FJ572210.1 | TBEV-Eur | Salem | - | 712 |
| NC_001672.1 | TBEV-Eur | Neudoerfl | - | 764 |
| U39292.1 | TBEV-Eur | Hypr | - | 458 |
| MK801805.1 | TBEV-Eur | Sipoo-8-Finland-2013 | - | 645 |
| LC171402.1 | TBEV-Eur | NL | W-Eur | 442 |
| MG243699.1 | TBEV-Eur | N5-17 | - | 412 |

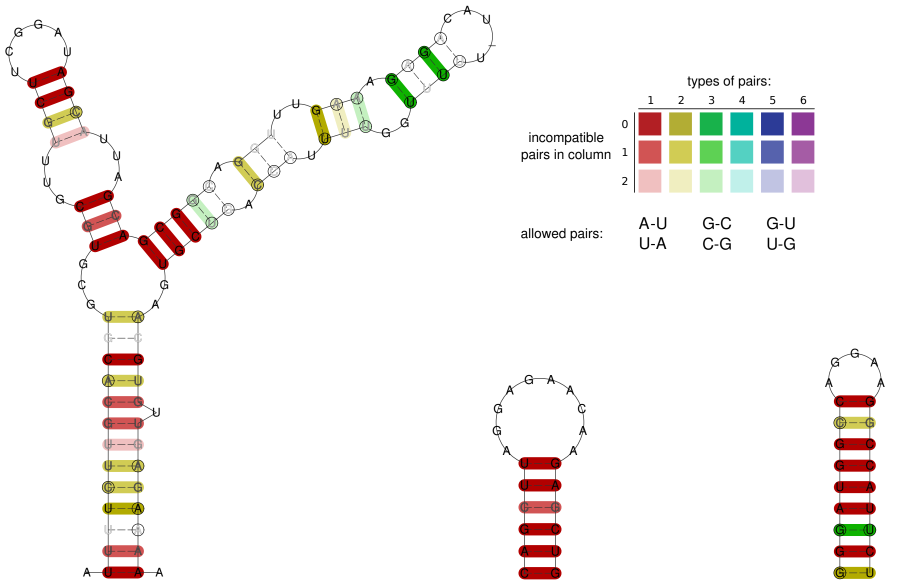


SLA CSA CSB

**Figure S1:** Consensus secondary structures of conserved RNA elements in the TBEV 5'UTRs. Base pair coloring indicates the level of covariation, following the default `RNAalifold` color scheme (top right). Circled nucleotides indicate compensatory mutations in the respective columns of the underlying multiple sequence alignments (MSAs). Structural MSAs for all elements shown here are available at the `viRNA` GitHub repository (`https://github.com/mtw/viRNA`).

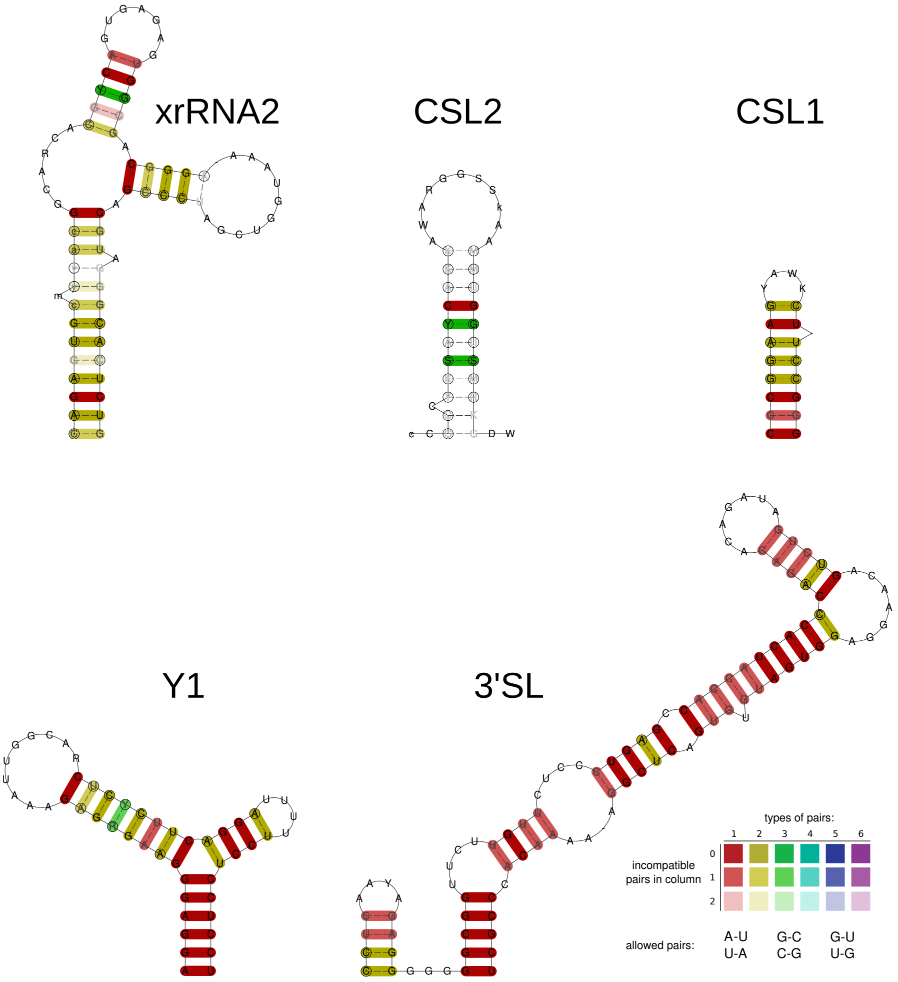


**Figure S2:** Consensus secondary structures of conserved RNA elements in the core region of TBEV 3'UTRs. See main text for details and Fig. S1 for a description of base pair coloring.

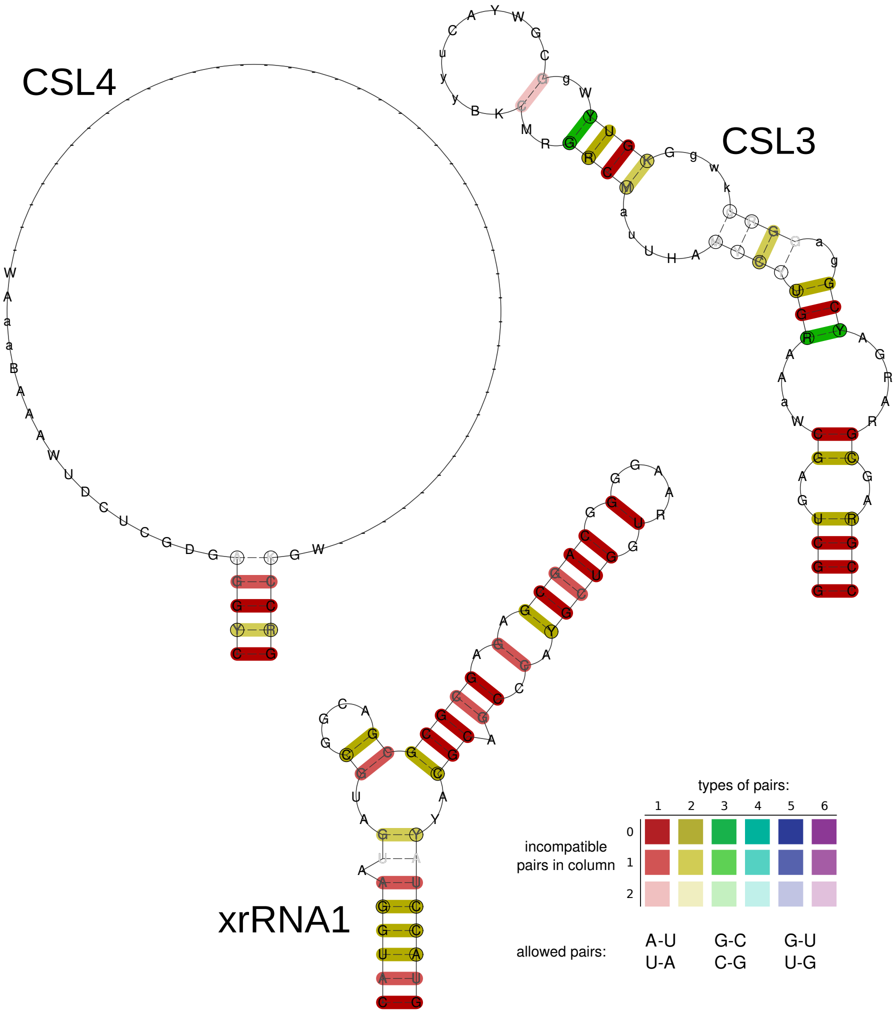


**Figure S3:** Consensus secondary structures of conserved RNA elements in the variable region of TBEV 3'UTRs. The large amount of gap characters in the hairpin loop of the CSL4 consensus structure is due to an insertion of a poly-A tract in one copy of the CSL4 element, observed in several TBFV-Eur strains. See main text for details and Fig. S1 for a description of base pair coloring.
